# Performance of a scalable extraction-free RNA-seq method

**DOI:** 10.1101/2021.01.22.427817

**Authors:** Shreya Ghimire, Carley G. Stewart, Andrew L. Thurman, Alejandro A. Pezzulo

## Abstract

RNA sequencing enables high-contents/high-complexity measurements in small molecule screens performed on biological samples. Whereas the costs of DNA sequencing and the complexity of RNA-seq library preparation and analysis have decreased consistently, RNA extraction remains a significant bottleneck for RNA-seq of hundreds of samples in parallel. Direct use of cell lysate for RNA-seq library prep is common in single cell RNA-seq but not in bulk RNA-seq protocols. Recently published protocols suggest that direct lysis is compatible with simplified RNA-seq library prep. Here, we evaluate the performance of a bulk RNA-seq library prep protocol optimized for analysis of many samples of adherent cultured cells in parallel. We combine a low-cost direct lysis buffer compatible with cDNA synthesis (“in-lysate cDNA synthesis”) with Smart-3SEQ and examine the effects of calmidazolium and fludrocortisone-induced perturbation of primary human dermal fibroblasts. We compared this method to normalized purified RNA inputs from matching samples followed by Smart-3SEQ or Illumina TruSeq library prep. Our results show that whereas variable RNA inputs for each sample in the in-lysate cDNA synthesis protocol result in variable sequencing depth, this had minimal effect on data quality, measurement of gene expression patterns, or generation of differentially expressed gene lists. We found that in-lysate cDNA synthesis combined with Smart-3SEQ RNA-seq library prep allows generation of high-quality data when compared to library prep with extracted RNA, or when compared to Illumina TruSeq. Our data show that small molecule screens using RNA-seq are feasible at low reagent and time costs.

## Introduction

RNA sequencing (RNA-seq) is a popular tool in modern biology that has expanded our understanding of the regulation and complexity of the transcriptome of many species. Primarily used for differential gene expression (DGE) analysis, other applications include transcript variants detection, alternative splicing, pathway analysis and exploring cellular heterogeneity and diversity in stem cell biology (1, 2). The typical RNA-seq workflow starts with RNA extraction from samples, followed by cDNA synthesis, adaptor ligation, size selection and high-throughput sequencing. The sequences are aligned to a reference genome and/or transcriptome to generate a genome wide transcription map which includes the transcriptional structure and/or expression profile for each gene (3).

RNA-seq workflows have become progressively more efficient over time. There are almost 100 distinct protocols derived from the standard RNA-seq approach (2). By studying the effects of many perturbations in parallel on cellular gene expression, efforts such as the Connectivity Map (4) have uncovered novel mechanisms of disease and therapeutic targets. However, significant bottlenecks impede adoption of highly parallelized methods in the absence of robotics equipment. Highly efficient RNA extraction is simple to perform in a small number of samples but becomes costly and laborious at the scale of hundreds of samples. Moreover, highly multiplexed RNA-seq library prep generally requires costly commercial kits.

Prior work has demonstrated that qPCR can be accurately performed using bulk cell samples without RNA extraction. These methods are generally based on the use of a lysis buffer compatible with cDNA synthesis and downstream sample processing and have been applied to single cell RNA-seq (5-9). Recently, Foley et al (10) developed a simple method (Smart-3SEQ) for rapid RNA-seq library prep which was tested with laser micro dissected tissue applied to cDNA synthesis reaction and with purified RNA, taking up to four hours on average and with low reagent costs.

We hypothesized that bulk RNA-seq can be performed in a multi-well plate format, and at low cost by performing RNA-seq library prep cDNA synthesis using in-lysate RNA followed by Smart-3SEQ. We performed an experiment comparing RNA-seq library prep using cDNA synthesis with in-lysate RNA versus RNA purified from the same samples. We used purified RNA for TruSeq RNA-seq library prep as a gold standard (Figure 1). Samples consisted of human dermal fibroblasts from 6 different human donors exposed to vehicle, calmidazolium (10 μM), or fludrocortisone (10 μM) for 6 hours. We examined the performance of RNA normalization-free in-lysate cDNA synthesis based on sequencing quality, sequencing depth, quantification accuracy and detection of differentially expressed genes (DEG) compared to purified RNA samples. We found that in-lysate cDNA synthesis allows generation of high-quality sequencing data without RNA input normalization. The gene expression profile of in-lysate RNA highly correlated with purified RNA. When compared to our gold standard, both methods showed similar performance for detection of DEGs. Thus, our data suggests that both methods tested are highly comparable to our gold standard for RNA-seq library prep.

**Figure 1:**
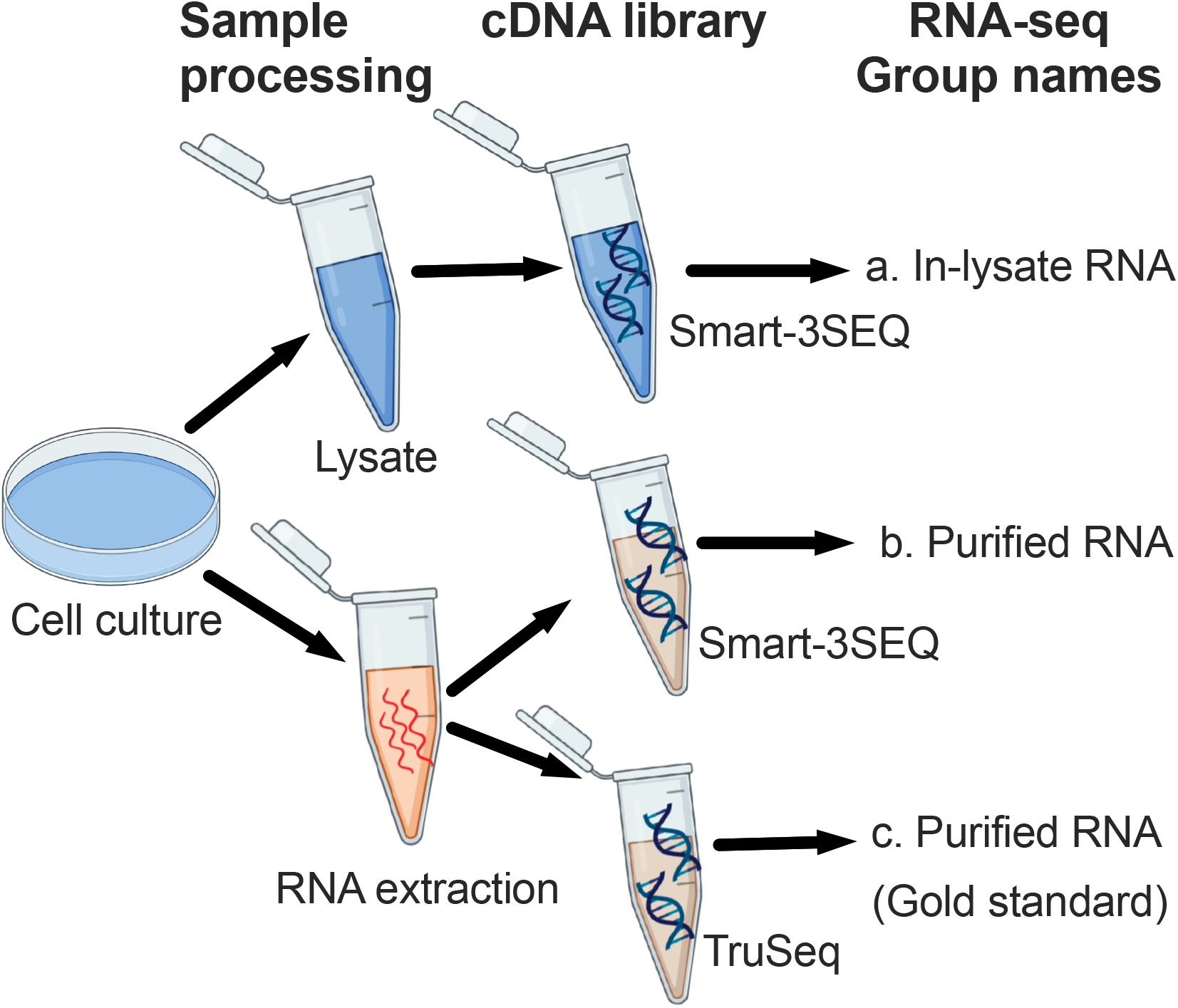
In-lysate RNA seq library prep performance: experimental overview. We used Smart-3SEQ to compare RNA-seq libraries prepared directly from lysate (a) and from extracted RNA (b). We used TruSeq libraries from matching RNA as the gold standard (c).18 samples(6 donors, 3 treatments) were used for each method.

## Results

### Effects of variable RNA mass loading on in-lysate RNA-seq library prep

Normalization of RNA input at scale is costly and laborious, so we examined the performance of normalization free in-lysate cDNA synthesis. We hypothesized that the number of alignable reads would be proportional to RNA concentration. As shown in Figure 2, in normalization-free samples (in-lysate RNA) the variable RNA concentration did not correlate with alignable reads (r^2^=0.145). As a comparison, we analyzed the number of alignable reads in libraries normalized for input RNA concentration (purified RNA). While the number of reads was on average 0.5 log higher than in-lysate RNA samples, there was similar variability in the number of reads in both approaches (log total counts 6.3 ± 0.4 vs 6.8 ± 0.3 mean/standard deviation for in-lysate vs purified RNA, respectively). These data suggest that in-lysate cDNA synthesis allows RNA-seq library prep without RNA input normalization. We then investigated the effects of in-lysate cDNA synthesis and RNA-seq library prep on sequencing data quality.

**Figure 2:**
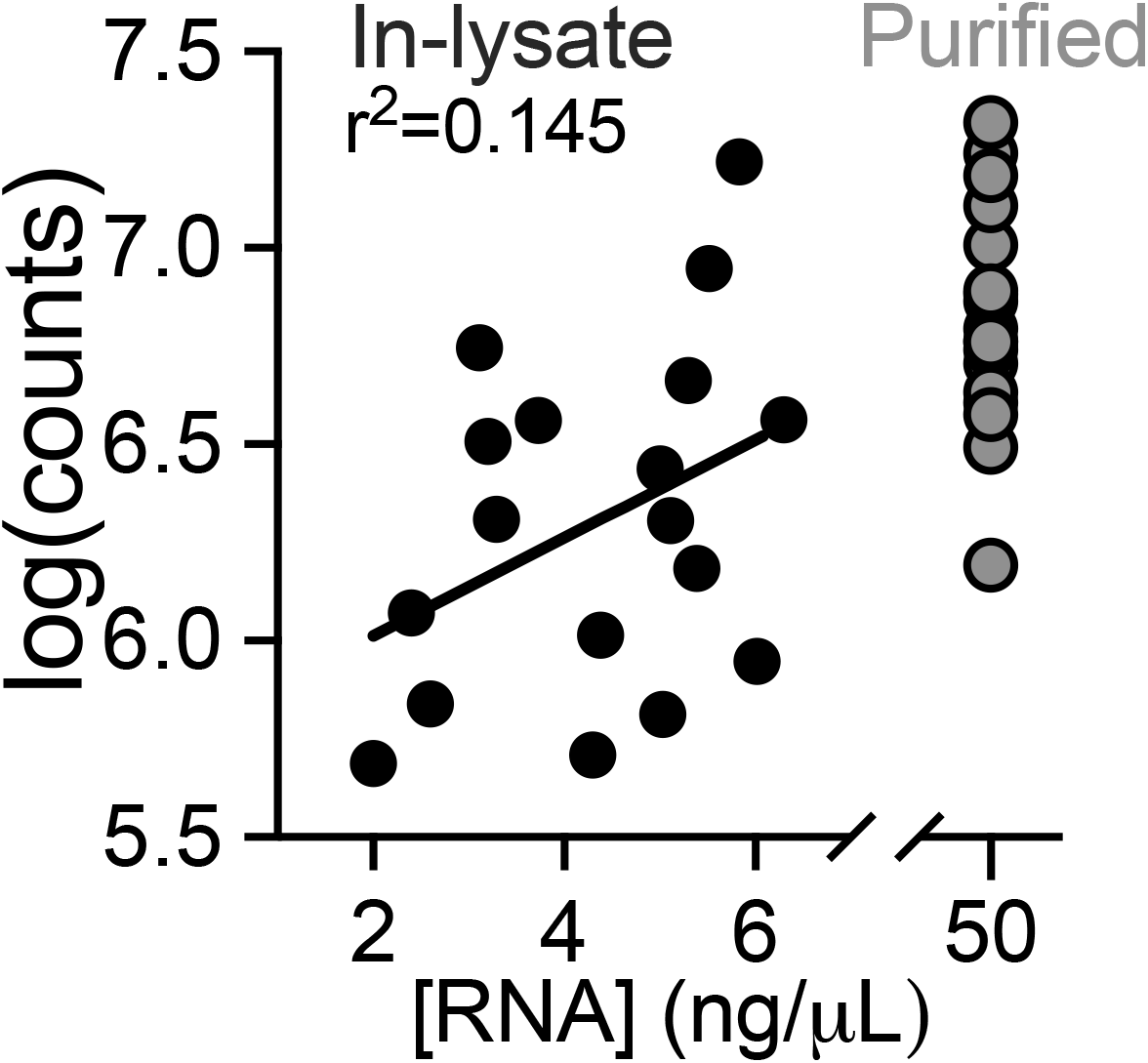
Effect of RNA loading without normalization on sequencing output. Scatter plot of total number of counts (y-axis) vs loaded RNA concentration(x-axis) for In-lysate library prep (In-lysate RNA). RNA concentration for Total RNA was normalized prior to cDNA library prep.

### In-lysate RNA-seq library prep does not negatively affect sequencing data quality

Quality control of RNA-seq data is critical for an accurate downstream data analysis. We hypothesized purified RNA samples would have better quality than in-lysate RNA samples due to their higher sequencing depth and likely lower presence of contaminating molecules during cDNA synthesis. We used FastQC (11), a quality control tool for high throughput sequencing data. Figure 3 shows average quality scores across each base position and the per sample distribution of quality scores. Both methods resulted in high-quality data with average Phred scores above 35. Remarkably, the variability in counts shown in Figure 2 for in-lysate RNA samples did not negatively affect the quality of RNA-seq data. Next, we investigated the effect of library prep methods on RNA-seq gene expression quantification accuracy.

**Figure 3:**
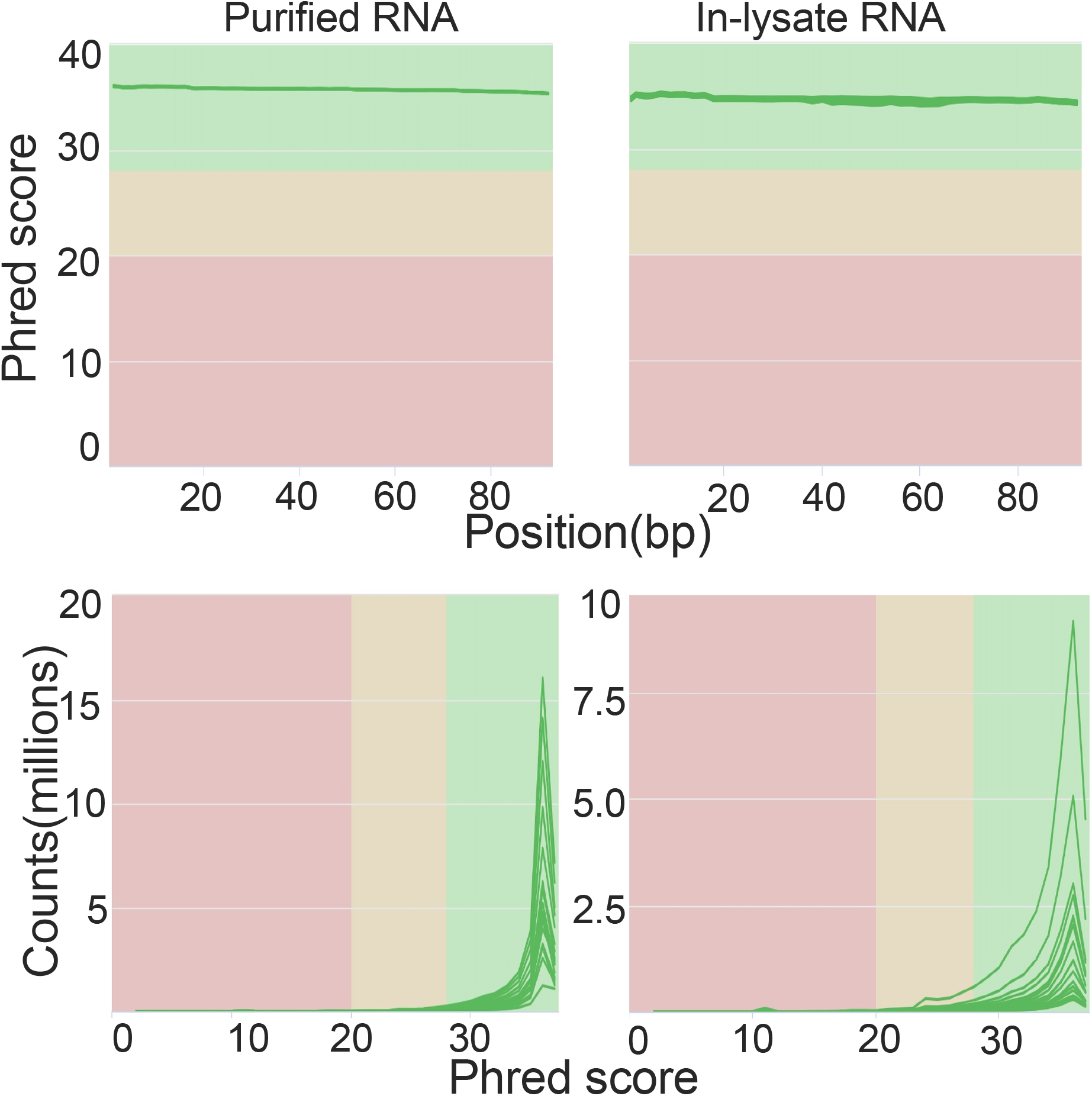
Quality control of sequencing data. Left panels corresponds to Purified RNA, right panels corresponds to In-lysate RNA (a) average quality scores (phred) across each base position in the reads. (b) per sample distribution of quality scores. Region labeling red= poor, yellow= acceptable, green= good.

### Gene expression profiles of in-lysate and purified RNA library prep highly correlate

Use of in-lysate RNA for RNA-seq library prep may result in unexpected biases in the type of mRNA molecules recovered that could either mask or accentuate variation between samples. We hypothesized that in-lysate and purified RNA library prep for RNA-seq would result in highly correlated data from matching samples. We randomly subsampled gene counts to equalize each set of matching samples in order to compare gene expression independent of sequencing depth. We then plotted the correlation within each treatment group: Vehicle, calmidazolium and fludrocortisone as shown in Figure 4. Out of three groups, vehicle showed highest correlation between methods, followed by Fludrocortisone and calmidazolium. Overall, all three treatment groups had high correlation between methods (R^2^ 0.96-0.99). These data show that gene expression profiles generated from RNA-seq libraries prepared with in-lysate versus purified RNA are highly comparable. Next, we used DESeq2 (12) to perform DEG analysis for data generated by each library prep method, and compared the list of detected genes.

**Figure 4:**
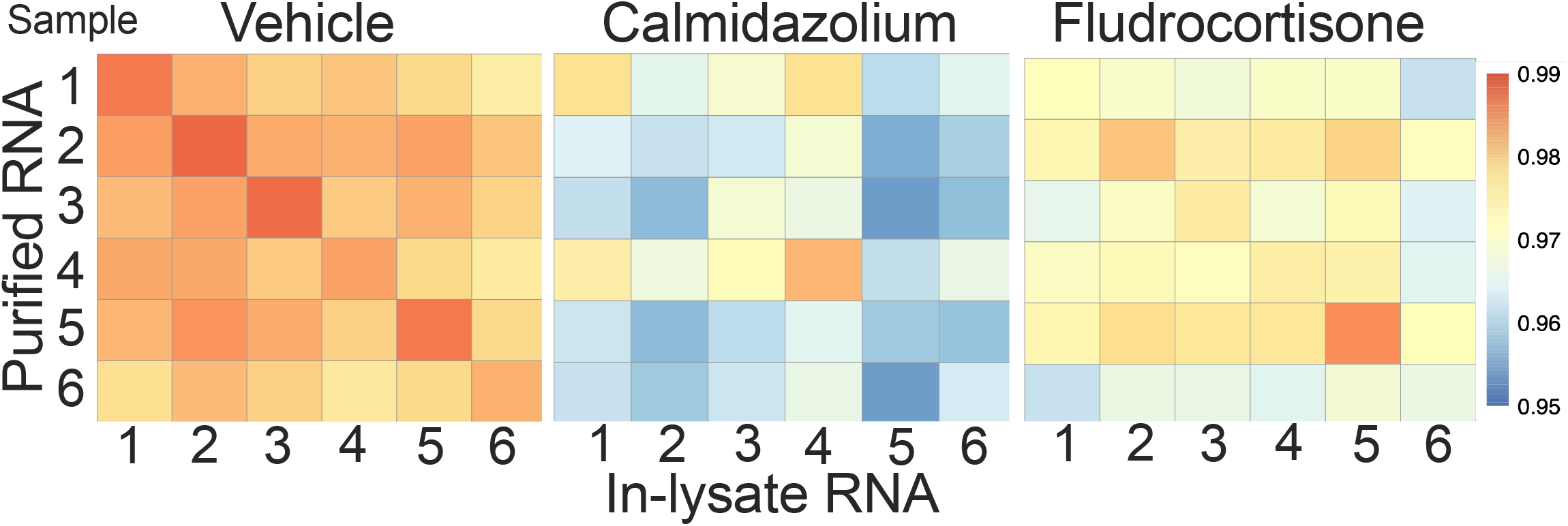
Gene expression correlation across methods. Gene expression correlation across methods for 3 treatment groups: Vehicle, Calmidazolium and Fludrocortisone. Depth normalized data

### DEG analysis reflects similar response patterns between in-lysate and purified RNA for default threshold

Variable sequencing depth can affect quantification of low-expressed genes for accurate differential expression analysis. We hypothesized that the normalized data would produce similar number of significant DEGs between methods for each treatment group.

Figure 5 shows MA plots for each method and treatment. DESeq2 uses the average expression strength of each gene across all samples as its filter and then removes all genes with mean normalized counts below a certain threshold, calculated using multiple tests. The chosen threshold, by default maximizes the number of genes found at a specified target FDR (10%). In Figure 5, the genes that contain enough information to yield a significant call at an estimated FDR of 10% are colored black. The calmidazolium treatment seems to induce more changes in gene expression compared to fludrocortisone treatment group (calmidazolium vs fludrocortisone, purified RNA: 1649 vs 639; in-lysate: 1059 vs 627). Overall, purified and in-lysate RNA samples reflects similar response patterns with default adjusted p-value within treatment groups. Next, we compared the significant genes using specific thresholds for log2 fold change (LFC) and adjusted p-value.

**Figure 5:**
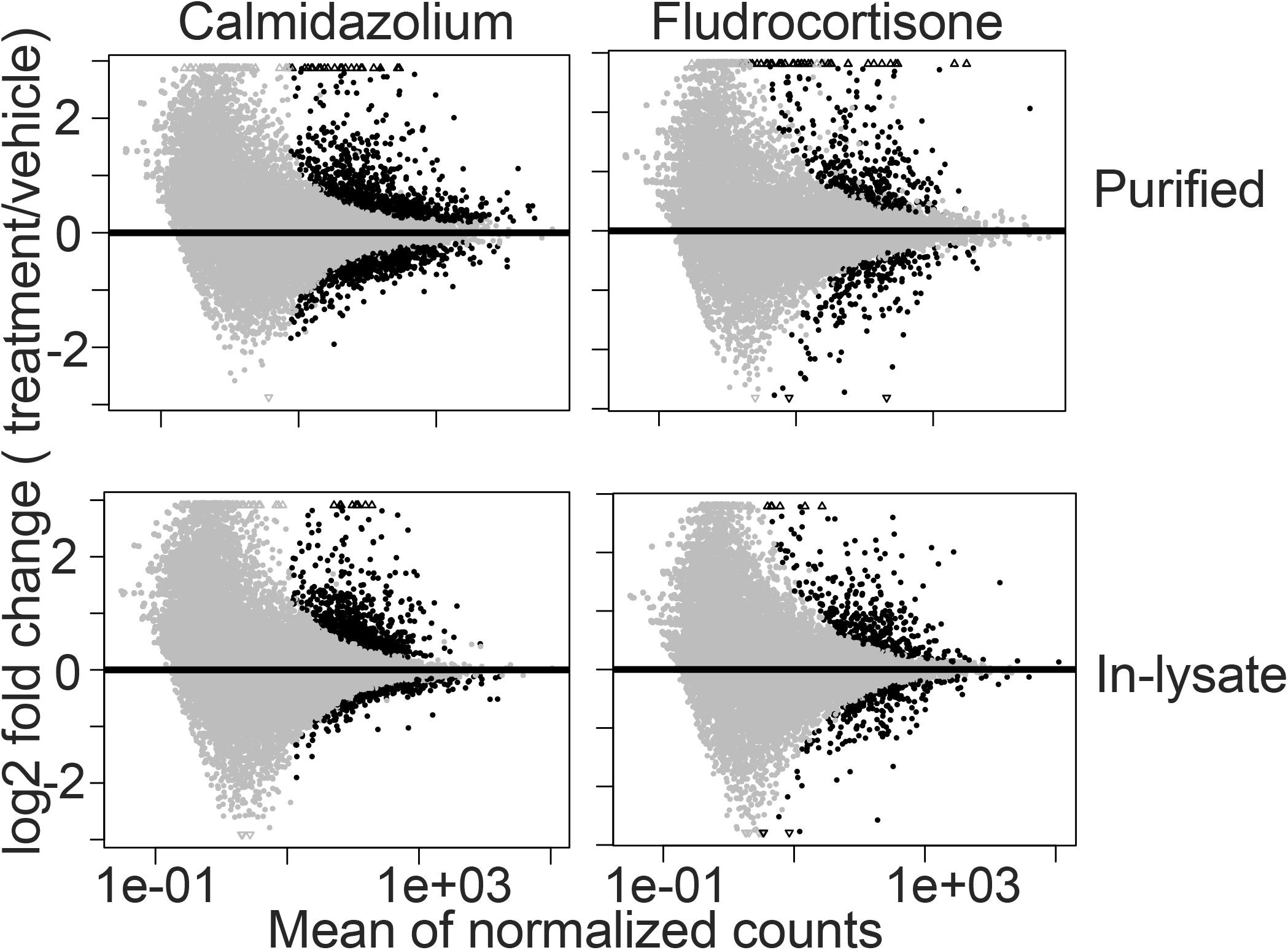
In-lysate RNA seq library prep and Purified RNA shows similar response patterns. Top panel: Purified RNA. Bottom panel: In-lysate RNA. Genes with adjusted p-value < 0.1 are labeled black. Small triangles at the top and bottom of plots indicate genes outside of the plotting window.

### RNA-seq library prep from in-lysate and purified RNA show similar gene response patterns for a specified threshold

We hypothesized that in-lysate and purified RNA would show similar gene response patterns within each treatment group. Figure 6 shows volcano plots in which, using cutoffs of LFC = 1 and adjusted p-value <0.05, both calmidazolium and Fludrocortisone induced comparatively more upregulated than downregulated genes in both methods. Purified RNA samples had more significant genes in both treatment groups compared to in-lysate RNA samples (purified RNA vs in-lysate RNA, calmidazolium: 263 vs 192; Fludrocortisone: 250 vs 150). Overall, both methods reflect similar differential gene expression patterns. Next, we examined the magnitude correlation of differentially expressed genes between two methods.

**Figure 6:**
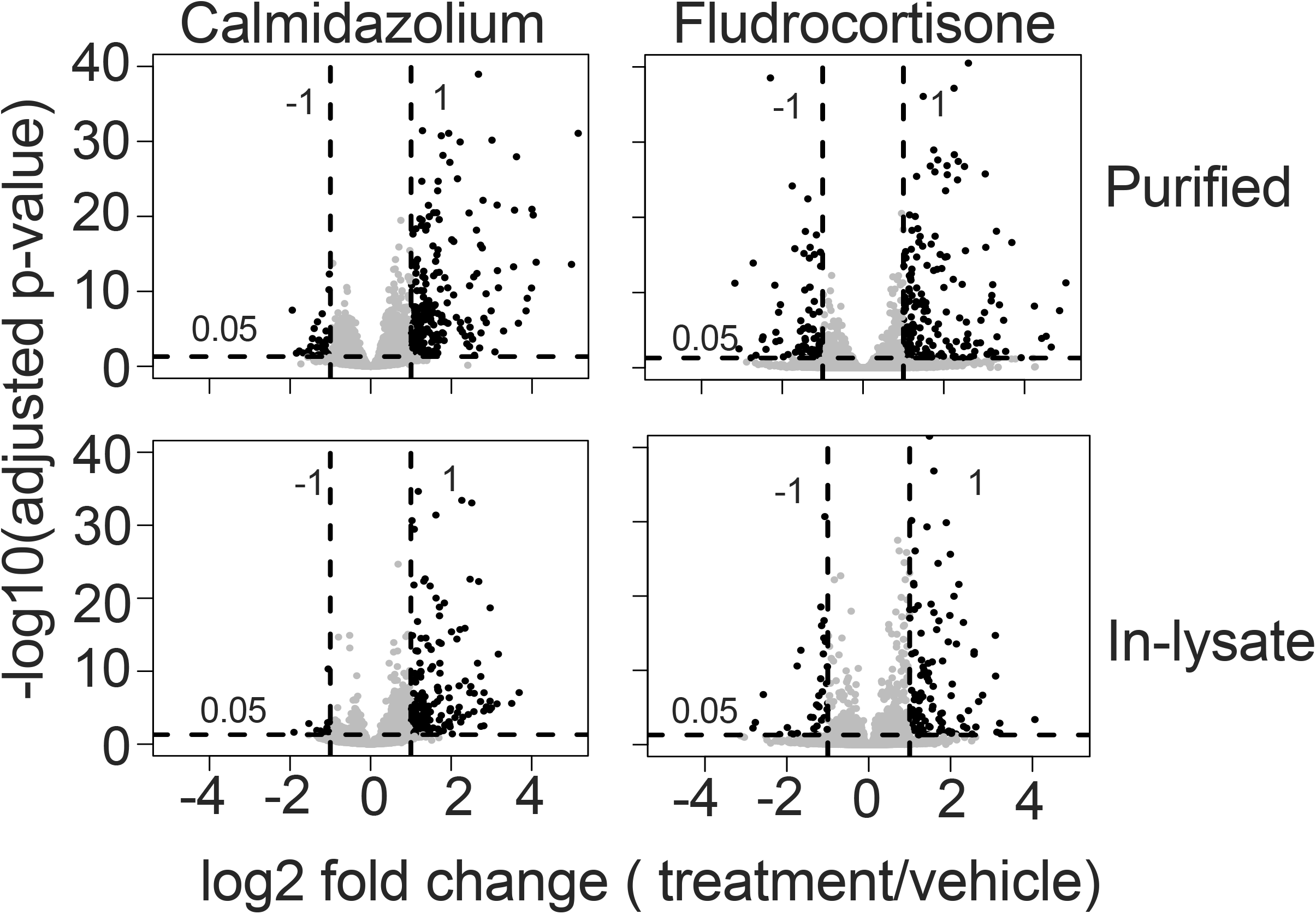
RNA seq library prep In-lysate vs from Purified RNA show similar gene response patterns. Volcano plots with cutoffs: log2FC +/-1, adjusted p-value 0.05

### The effect sizes of significant DEGs from in-lysate and purified RNA highly correlate with each other

Using the LFC and adjusted p-values, we generated “F-F plots” to further examine the relationship between DEGs from the different methods tested. We hypothesized that DEG LFC from purified and in-lysate RNA samples would highly correlate with each other. As shown in Figure 7, purified and in-lysate RNA genes showed positive linear correlation particularly for significant genes (adjusted p-value < 0.05). R^2^ for threshold-passing genes was 0.906 and 0.969 for calmidazolium and fludrocortisone treatment groups, respectively. This suggests that there is a high positive correlation between the LFC of genes that are significant in both methods. Next, we examined this concordance with respect to our gold standard (TruSeq) using rank-based and binary classification methods.

**Figure 7:**
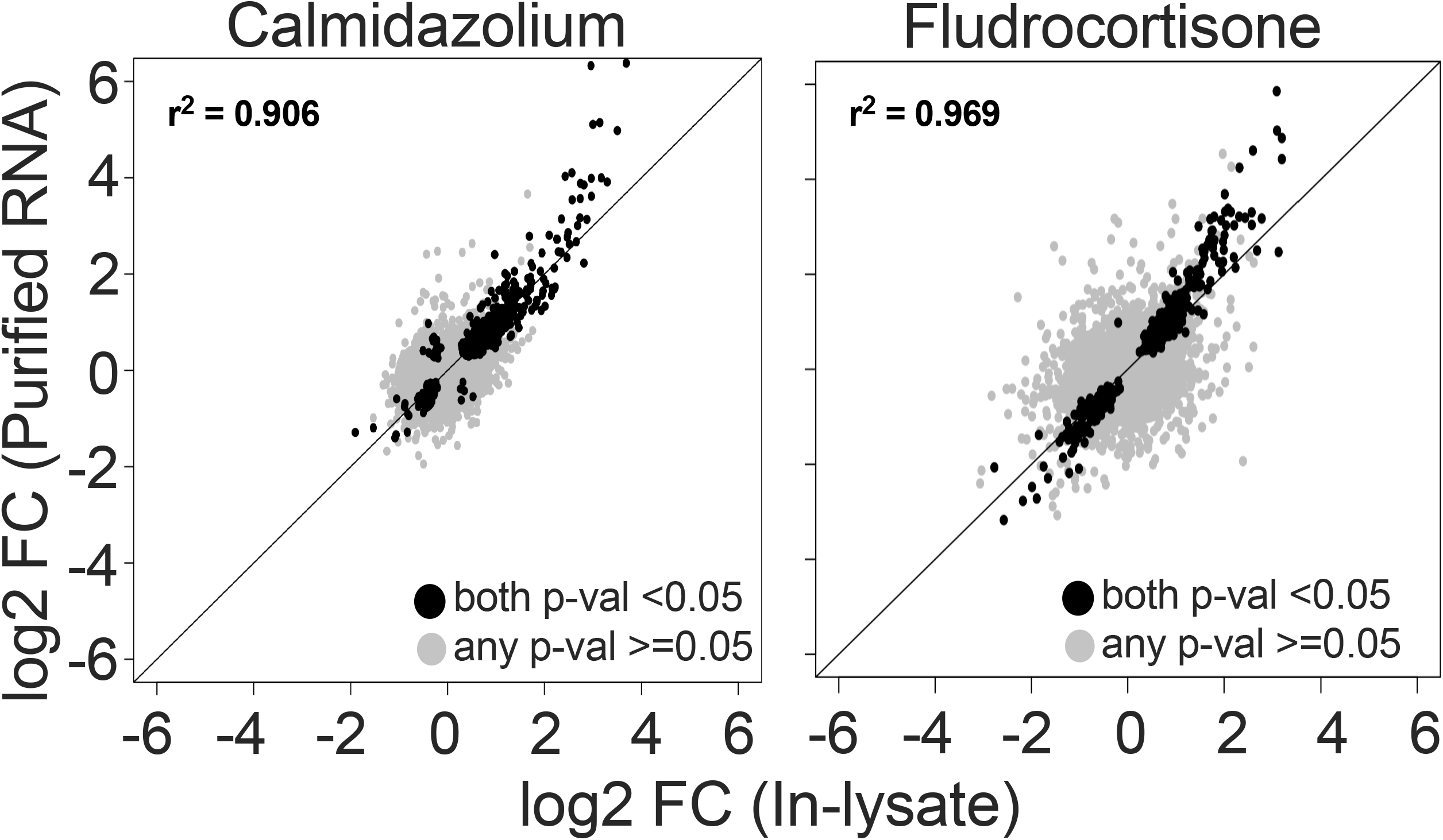
Purified and In-lysate RNA genes show positive linear correlation. F-F plot: log2 fold change of Purified vs In-lysate RNA. Genes with p-value < 0.05 in both methods are colored black. Grey genes indicate genes with higher p-value (>=0.05) in either of the two methods. R^2^ for black genes are shown in the figure.

### In-lysate and purified RNA show similar performance for calling DEGs

To further investigate how the gene expression pattern from in-lysate RNA and purified RNA library preps compare in performance, we plotted Receiver Operating Curve (ROC) and Precision-Recall (PR) curves for each treatment group (Figure 8), using the TruSeq dataset as a gold standard. We included PR curves in addition to ROC because of the high imbalance in our data for the positive (DEGs) and negative (non-DEGs) classes. To produce these curves, we used different cutoffs for adjusted p-values between zero and one, and the “true” class labels were genes with an adjusted p-value 0.05 for TruSeq. We hypothesized that purified RNA would perform better than in-lysate RNA when compared to our gold standard. In Figure 8, ROC and PR curves show that both methods performed better for the fludrocortisone treatment group compared to calmidazolium. Overall, purified RNA libraries performed slightly better than in-lysate RNA libraries for each treatment group and classification method. These data suggest that in-lysate RNA-seq library prep has a slight negative effect on performance of DEG analysis.

**Figure 8:**
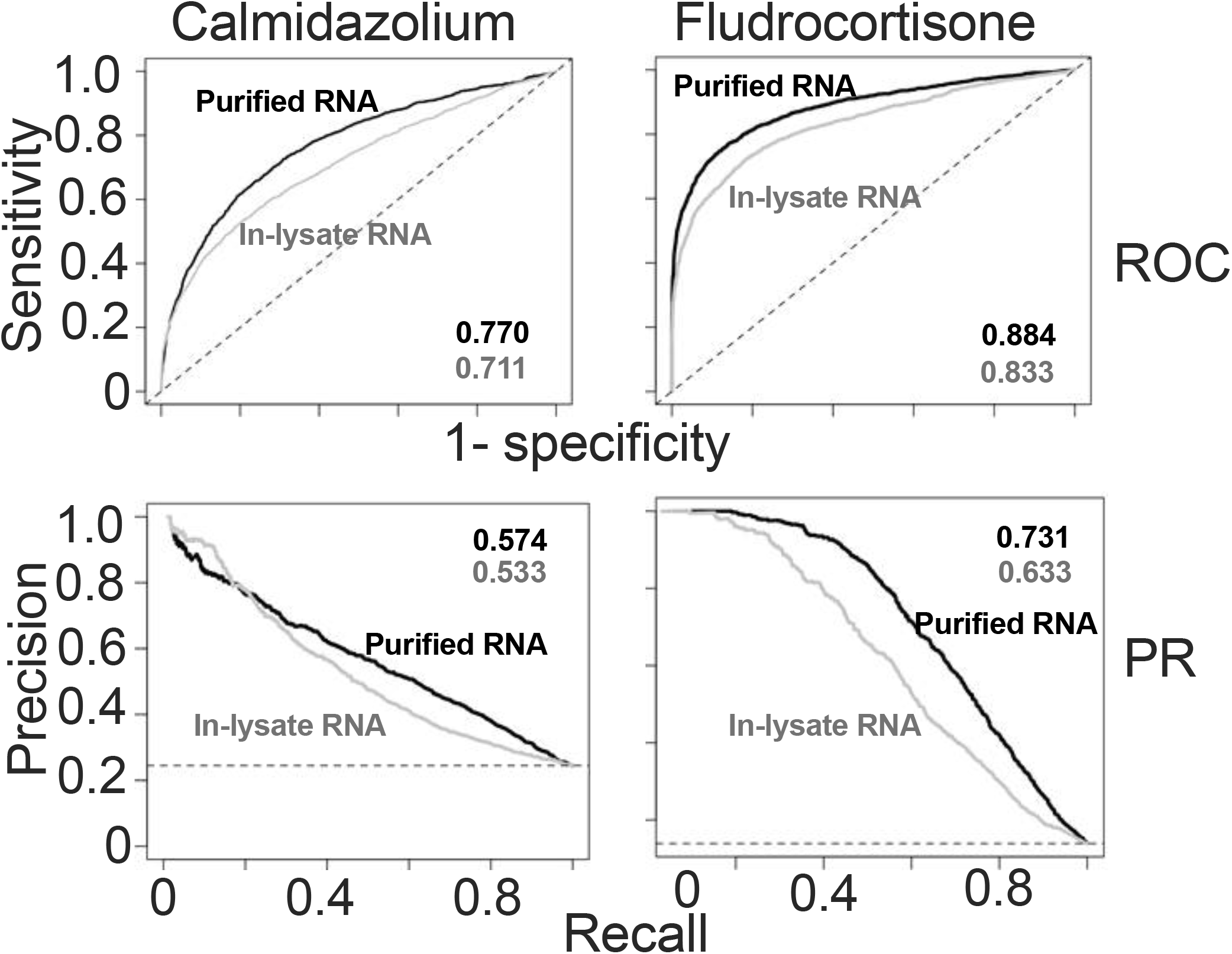
Method performance for DE genes calling compared to Illumina TruSeq (gold standard). Top panel: Receiver Operating Curve (ROC), Bottom panel: Precision-Recall cuve (PR). Area under the curve (AUC) indicated in figure.

### Similar DEG sets are called with in-lysate and purified RNA-seq library prep

One of the main goals of DEG analysis is to prioritize lists of genes highly likely to be differentially expressed between two conditions for subsequent pathway analysis. DEG lists generated with arbitrary FC and p-value cutoffs can be affected by methods used for RNA-seq library prep or data analysis, variable depths or reference transcriptomes. In contrast, rank-based methods allow easier comparisons between lists by focusing on genes/transcripts enriched near the top of the rankings (13-17). We hypothesized that data from in-lysate RNA and purified RNA library preps ranked by adjusted p-value and absolute fold change would highly correlate near the top of the ranked list when compared to data from TruSeq libraries as a gold standard (Figure 9). We found that there was a high (70-75%) overlap in lists of DEGs when the threshold used was between 300-500 top DEGs. This concordance dropped quickly prior to rising again as expected due to random matching. This data suggests that between 300-500 genes are likely to be modulated by fludrocortisone in human dermal fibroblasts. Moreover, these data suggest that both methods produce ranked lists that are highly comparable to our gold standard for RNA-seq library prep, and would yield similar results in downstream functional analysis of DEG lists.

**Figure 9.**
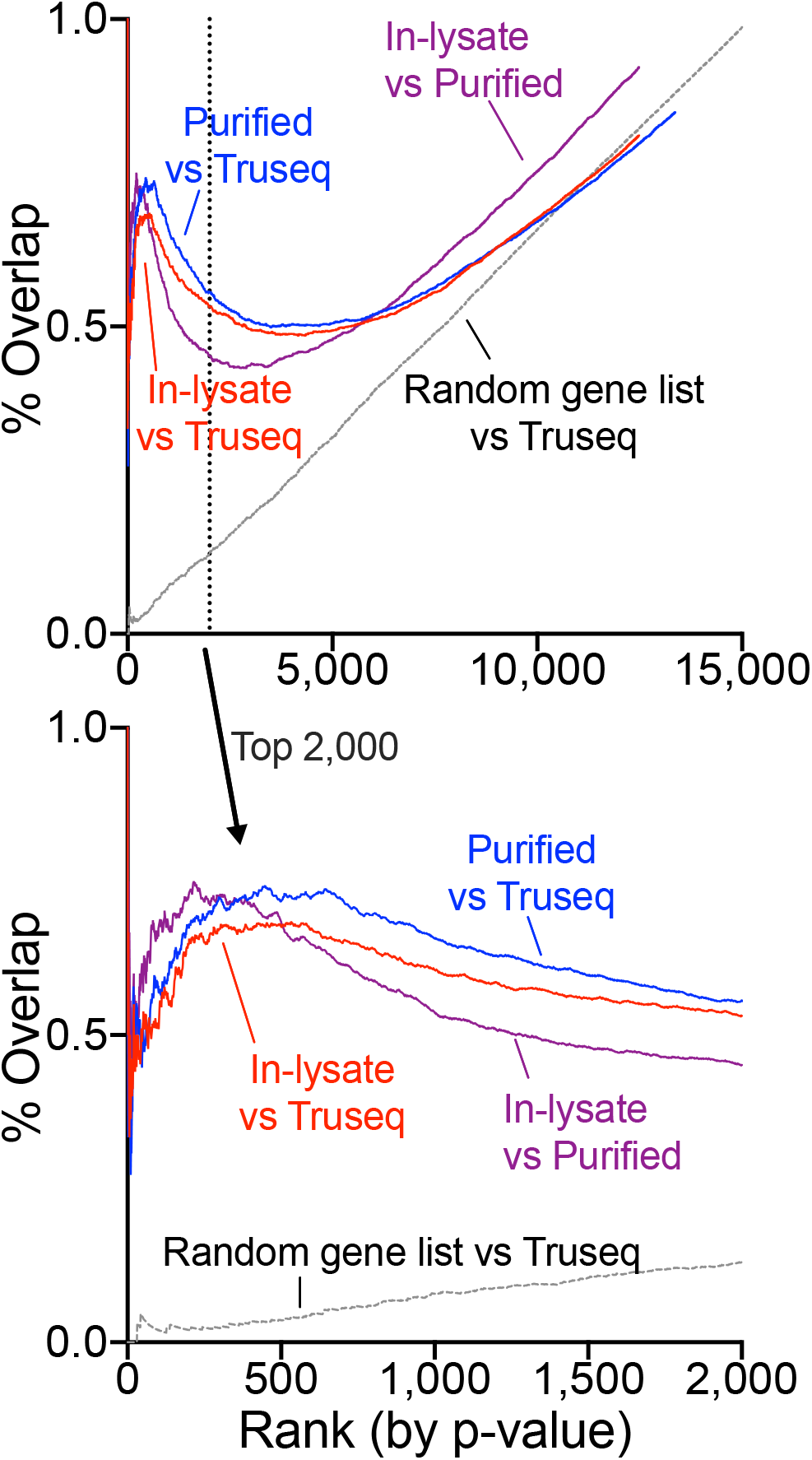
Differentially expressed gene-calling concordance between methods. Differential gene expression lists for libraries prepared with Smart-3’seq (In-lysate RNA vs purified RNA) and Illumina Truseq (extracted RNA alone, gold-standard) were sorted by p-value. For each rank position, the % overlap between lists was calculated. A randomly re-ranked list for all detected genes was compared to the Truseq data. **Top panel shows all gene ranks, bottom panel shows only top 2000**

## Discussion

Recent methods for transcriptome analysis on single cells show that RNA extraction can be skipped by using lysis buffer that is compatible with cDNA synthesis and downstream analysis. These studies show that direct lysis gives equivalent or superior RNA recovery than commercial column or magnetic beads-based RNA extraction methods which makes this method rapid, cheap and highly effective (5). Here, we expand this concept to library prep using samples of adherent cells growing in multi-well plates, which is a very common high throughput assay model.

We demonstrate that high quality bulk RNA-seq can be performed without RNA isolation for library prep. Moreover, when combined with methods for simplified multiplex library prep using off the shelf reagents, significant time and reagent cost savings are achieved. We chose to use Smart-3SEQ library prep because it has fewer steps and uses lower volume of common reagents compared to many other RNA-seq library prep methods. It allows accurate quantification of transcript abundance with low RNA input making this approach rapid, robust and cost-efficient (10). Performing in-lysate Smart-3SEQ library prep, we have shown the success of our approach on: 1) all quality control criteria were achieved, and 2) gene expression profiles, DEG analysis, and response patterns reported by in-lysate and purified RNA library highly correlate with each other.

Thus, the approach presented here rapid, robust and highly cost-efficient RNA-seq of adherent cell samples in parallel. The reagent costs for generating lysates used for library prep are negligible compared RNA extraction using silica columns or magnetic beads. Moreover, this approach takes only approximately 4-6 hours from sample processing to final sequencing-ready library. However, there are some caveats to the approach presented here. The amount of RNA loaded for in-lysate cDNA synthesis was not normalized which resulted in variable and low-sequencing depth across samples. Although this issue didn’t negatively affect the quality or the gene expression profile of the samples, we did get lower numbers of DEGs from in-lysate RNA samples. Moreover, Smart-3SEQ sequences only the 3’ fragment with Poly(A) tail of each transcript and is not sensitive to transcript length. Although this allows for a simple and accurate quantification of transcript abundance, information regarding splicing or genotypes cannot be detected unless the splicing junction is near the 3’ end of the transcript (10). We used a matching dataset generated with libraries prepared from extracted RNA and processed with the Illumina TruSeq as a gold standard. Whereas a digital gene expression method such as Smart-3SEQ is not expected to match the data generated with a full-length transcript library prep method like TruSeq, it is a useful gold-standard as it’s frequently used and allows further head-to-head comparisons between our test groups. We detected a decrease in within-sample correlation between methods upon small-molecule perturbation compared to vehicle-treated samples. This effect was more marked with calmidazolium treatment. We speculate various possibilities: 1) the expression profile of human fibroblasts may become more similar between individuals upon stimulation, 2) in-lysate RNA library prep may capture mRNA exported to the cytoplasm with may occur differentially between individuals, or 3) perturbation with calmidazolium may cause changes undetected by our methods in the ability to correctly fragment and process mRNA.

Other transcriptional profiling platforms have been developed that facilitate processing of hundreds of samples in parallel. Luminex L1000, used for the Connectivity Map (4) is a cost-effective platform for transcriptional profiling; however, it only allows the measurement of approximately 1000 genes making this approach less efficient and more susceptible to signal-noise problems (18). Moreover, the inference method applied to estimate expression of the remaining transcriptome may not be accurate for specific sample types. Pooled library amplification for transcriptome expression (PLATE-Seq) is another low-cost approach for genome wide profiling which highly compares with other large-scale profiling, however it is based on RNA purification (4). Another approach, Digital RNA with perturbation of Genes (DRUG-seq) skips RNA extraction by using direct lysis approach and cuts down the library prep time and cost. Here, we chose SMART-3SEQ as it relies on enzymatic fragmentation of RNA rather than tagmentation of pooled libraries which is costly and can be variable. The approach presented here is therefore highly compatible with high throughput drug screening at a low cost.

Highly-parallel assessment of perturbation-transcriptome responses in biological samples can be an unbiased and data-rich approach to screen compounds and to test hypothesized mechanisms of disease. Other novel approaches such as single cell RNA-seq-based genetic perturbation analysis together with small-molecule-based approaches as presented here provide a nimble toolbox to uncover novel biology.

## Methods

### Cell culture

Human primary dermal fibroblasts (previously obtained with approval from the University of Iowa Intitutional Review Board, IRB# 201311783) were obtained using the method described in (19). Cells were maintained in DMEM (Gibco, Dublin, Ireland, Item No. 11965-092) with 20% FBS (Gibco, Dublin, Ireland, Item No. 26140-079) and Pen/Strep 1:100 (Gibco, Dublin, Ireland, Item No. 15140-122) and gentamycin 25μg/mL (IBI Scientific, Dubuque, IA, Item No. IB02030), at 37°C with 5% CO_2_ and were used at passage #4, near confluency, in 24 well plates. Samples were treated for 6 hours as indicated with 10μM Calmidazolium chloride (Cayman Chemical Company, Ann Arbor, MI, Item No. 14442) or Fludrocortisone acetate (Sigma-Aldrich, St. Louis, MO, Item No. F6127).

### RNA preparation

Total RNA was extracted and purified from cell cultures using RNeasy plus mini kit from QIAGEN (Hilden, Germany). In-lysate RNA was prepared by washing cell cultures PBS, followed by with a buffer adapted from (5-9) consisting of 0.1% (1mg/mL) BSA (New England Biolabs, Ipswich, MA, Item No. B9000S) / 0.3% Igepal CA 630 (Sigma-Aldrich, St. Louis, MO, Item No. I8896) for 5 minutes while on ice. Based on these studies, we estimated an appropriate cell/buffer volume ratio to be between 100-500 cells per μL buffer for adherent cells, an excessively high ratio may negative affect compatibility with cDNA synthesis. For our cell culture vessel (24 well plate), near confluent cultures correspond to an average 100,000 cells/well. We chose a ratio 200 cells/μL buffer, so 500ul of Igepal/BSA buffer was used per well. The lysate was aspirated and was only mixed with the pipette tip once in a collection tube. Total RNA concentrations were measured using Qubit HS RNA and Nanodrop (Thermo Fischer Scientific, Waltham, MA). Samples were stored at -80°C until library prep.

### Library preparation

Extracted RNA and cell lysate (above) was used for preparing sequencing libraries according to the Smart-3SEQ protocol (10). Illumina TruSeq libraries were prepared at the University of Iowa Genomics Facility using standard Illumina TruSeq stranded mRNA protocols. Concentration and fragment size of prepared library was measured using Qubit HS and bioanalyzer. Fragment size for Smart-3SEQ library was ∼500bp which falls within the recommended 200-600bp.

### Sequencing

Smart-3SEQ cDNA libraries were prepared with a 40% spike-in of phiX and sequenced on Illumina NovaSeq 6000 SP Flowcell instrument (approximately 400M PF Reads per lane) configured for 100nt read 1 and 8nt for the P7 index read. TruSeq mRNA-seq libraries were sequenced on a single lane of an Illumina HiSeq 4000 in 75PE configuration.

### Data processing

The base calls were demultiplexed and converted to FASTQ format using Illumina bcl2fastq. For Smart-3SEQ data, adapters were trimmed, UMI sequence and G-overhang were extracted and polyA tails were removed according to the suggested pipeline in data processing protocol (10).

Quality control analysis was done with FastQC and MultiQC (20). The processed fastq files were aligned using HISAT2(21) and hg38 reference genome. Aligned transcripts were counted by featureCounts from the Subread package excluding duplicate reads (22). Since only one sequencing fragment per RNA molecule is generated using Smart-3SEQ method, normalization isn’t necessary as read counts are directly proportional to the transcript abundance(10). Reads for TruSeq mRNA-seq were counted equally except for the following unnecessary steps: preprocessing; adapter trimming, G overhang and PolyA tail removal. Normalization by transcript length was done to obtain transcript abundance.

### Gene expression analysis

Read counts were used as input for differential gene expression analysis by DESeq2 (12). Default options were used. Correlation and F-F plot was created using R function. ROC and PR curve were plotted using ROCR package(23).

### Data availability

The data used in this study is available in the NCBI Gene Expression Omnibus, accession GSE164650

## Acknowledgements

We thank Einat Snir and Kevin Knudtson at the University of Iowa Genomics Division and Phil Karp, Ping Tan, and Shu Wu at the University of Iowa Tissue and Cell Culture Core for significant technical support during adoption of the methods described here.

